# Conditional replacement of the mouse LH receptor with GFP, enabling imaging of cell migration during ovulation

**DOI:** 10.64898/2026.05.21.726840

**Authors:** Corie M. Owen, Katie M. Lowther, Deborah Kaback, Laurinda A. Jaffe, Siu-Pok Yee

**Affiliations:** Department of Cell Biology, University of Connecticut Health Center, Farmington, CT 06030 USA; Center for Mouse Genome Modification, University of Connecticut Health Center, Farmington, CT 06030 USA

**Keywords:** luteinizing hormone receptor, ovary, mouse, ovulation, granulosa cell migration

## Abstract

To facilitate the investigation of signaling by the luteinizing hormone receptor (LHR), we created a mouse line called *Lhr*-COIN. This line allows for the conditional replacement of the *Lhr* coding sequence with enhanced green fluorescent protein (eGFP), resulting in both a conditional knockout line and a reporter line. By breeding these mice with mice expressing Cre recombinase, we generated mice in which either one or both *Lhr* alleles were replaced with eGFP. Notably, mice in which one *Lhr* allele in the granulosa cells was replaced with eGFP exhibited normal LH responsiveness. This enabled live imaging of LH-induced migration of LH-receptor-expressing granulosa cells within preovulatory ovarian follicles. The *Lhr*-COIN mouse line holds significant potential for future research on LHR function and localization in the ovary and other tissues.

## Introduction

Luteinizing hormone (LH) acts by way of a G-protein coupled receptor that controls many essential functions in ovary^1^ and testis^2,3^ of mammals and other vertebrates^4^. In mammalian ovaries, the luteinizing hormone receptor (LHR) acts to stimulate the progression of meiosis in oocytes within preovulatory follicles, by signaling in the mural granulosa cells^5^. Activation of the LHR also stimulates ovulation^6–8^ and the formation and function of the corpus luteum that supports pregnancy^9^. In addition, the LHR is expressed in the theca cells of preovulatory and smaller follicles, and in ovarian interstitial cells (see^10^), although its role in these other cells is less well defined. In the testis, the LHR is expressed in Leydig cells, where it is essential for testosterone production, cell proliferation and differentiation, development of the male reproductive tract, and spermatogenesis^2,3,11,12^.

To facilitate studies of LHR localization and function, we previously generated a mouse line in which the endogenous LHR was tagged with a 9-amino acid hemagglutinin epitope on the N-terminus (HA-*Lhr*), which allowed us to visualize the localization of LHR protein in fixed immunolabelled sections of ovary^10^. Analysis of ovaries fixed after inducing release of endogenous LH showed that LH stimulates LHR-expressing granulosa cells to migrate inwards within preovulatory follicles^13^. LH stimulation of granulosa cell migratory behavior has also been demonstrated using isolated granulosa cells, by measuring their movement across a porous membrane^14^. Granulosa cell migration may have a functional role in the ovulatory process, but this remains to be tested.

Here, we generated a mouse line (*Lhr*-COIN), in which the *Lhr* coding sequence is conditionally replaced in-frame with exon 1 by eGFP, using a conditional-by-inversion strategy. By breeding the *Lhr*-COIN mice with mice expressing Cre recombinase, we obtained mice with cells that express one copy of the LHR and one copy of eGFP. One copy of the LHR was sufficient for normal function, and the co-expression of eGFP allowed visualization of the LHR-expressing cells. We used these mice to visualize the LH-induced migration of LHR-expressing granulosa cells in live preovulatory follicles. More generally, this mouse line will be useful for investigating the localization of the LHR in the testis^2,3^ and in other tissues where LHR expression has been reported^15–18^.

In addition to its utility as an LHR reporter, the *Lhr*-COIN mouse line also allows conditional disruption of both *Lhr* alleles in specific cell types, providing a conditional *Lhr* knockout. Two mouse lines with global deletion of *Lhr* have been generated previously^11,12^ and have been used to elucidate many aspects of LH signaling^15,19^. The conditional knockout line described here will be useful for testing the expression and function the LHR in particular cell types within the ovary, testis, and other tissues.

### Mice

The generation of the *Lhr*-COIN mouse line is described in the Results and in the Supplementary Methods^20^. This mouse line has been deposited at the Mutant Mouse Resource and Research Center at The Jackson Laboratory (Bar Harbor, ME) (MMRRC #076250, C57BL/6J-*Lhcgr*^*em2Laj*^/Mmjax, JAX stock #041561).

Mouse lines expressing Cre recombinase were obtained from The Jackson Laboratory and were bred to C57BL/6J mice in our lab. The *Hprt*^*Cre*^ line^21^ is JAX stock #004302, and the *Aro*^*Cre*^ line^22^ is JAX stock #027038.

Animal studies were performed in accordance with the Guide for the Care and Use of Laboratory Animals (National Academy of Sciences 1996) and were approved by the Institutional Animal Care and Use Committee at the University of Connecticut Health Center.

### Culture and imaging of isolated follicles

Large antral follicles (290-360 µm in diameter) were dissected from 23-26-day old mice, using fine forceps (Fine Science Tools, #11200-14). The follicles were placed in individual drops on 30 mm optically clear Millicell culture plate inserts (MilliporeSigma, PICM0RG50) as previously described^23^. The follicle culture medium^24^ contained 1 nM ovine follicle stimulating hormone (FSH) (National Hormone and Peptide Program, AFP7558C; currently available from Golden West Companies, Temecula CA), except as indicated. LH (National Hormone and Peptide Program, ovine LH-26; currently available from Golden West Companies) was introduced by moving the Millicell into a new dish containing 2.0 ml of medium with 10 nM LH.

For evaluation of the occurrence of GVB and ovulation, live follicles were imaged on 30 mm Millicells, using an upright microscope with a 20x/0.4 NA/~11 mm working distance LD Achroplan objective (Zeiss #44 08 44) and photographed using an iPhone 13 Pro camera attached with a LabCam microscope adapter (iDu Optics).

For evaluation of FSH-induced eGFP expression, the follicles were dissected as described above, except that the follicle culture medium contained no FSH. Follicles were placed on 12 mm optically clear Millicell culture plate inserts (MilliporeSigma, PICM01250) from which the 1 mm tall plastic feet had been cut off before mounting in wells with a glass coverslip base (Ibidi µ-Slide 2-well slide, #80287) using 360 µm adhesive spacers (Grace Bio-Labs, #620003). Each well contained 1.4 ml of medium without FSH. Follicles were incubated on the Millicell for ~3 hours and then imaged using a confocal microscope (LSM 980, Carl Zeiss Microscopy) with a 10x/0.5 NA/1.7 mm working distance Fluar objective (Zeiss #420140-9901-000) and an Airyscan detector. After imaging, the culture medium was removed from each well and replaced with 1.4 ml of medium containing 1 nM FSH. 23 hours later, follicles were imaged again using the same laser intensity and detector gain.

### Time lapse imaging of LH-induced granulosa cell migration in clusters of preovulatory follicles

For live imaging of granulosa cell migration, ovaries were isolated from 23-26 day old mice expressing one copy of *Lhr* and one copy of eGFP. Follicle groups containing 2-4 follicles, each 290-360 µm in diameter, were dissected from the ovary and cultured for 24 hours on 12 mm Millicell culture plate inserts mounted in wells as described above, except that 240 µm spacers were used (Grace Bio-Labs, #620002). The medium contained 1 nM FSH.

Time lapse imaging was conducted using a confocal microscope as described above, with a stage top incubator to maintain 37^°^C, 5% CO_2_, and ~100% humidity^25^. After an ~1 hour equilibration period in the stage top incubator, a pre-perfusion image was taken as the baseline. About 30 minutes later, 10 nM LH was introduced by removing all medium from the dish, and replacing it with 1.4 ml of medium containing LH. The microscope was then refocused on the clusters, and imaging was started 3-5 minutes after LH addition. Images were collected every 15 minutes for 20 hours. Images were reduced in scale by 50%, then saved as an AVI with JPEG compression and a frame rate of 15 frames per second.

To quantify the LH-induced relocation of eGFP fluorescence within the mural granulosa compartment, we defined 2 concentric regions: the total mural region, extending 100 µm inwards from the basal lamina, and an inner region that started 50 µm from the basal lamina and extended 50 µm inwards. Before perfusion, immediately after, and for every hour for 8 total hours, we measured fluorescence intensity in each region using Fiji^26^(https://pubmed.ncbi.nlm.nih.gov/22743772/). Intensity values were corrected for minor autofluorescence using measurements from wildtype follicles imaged under identical conditions. We then calculated the percentage of the total eGFP fluorescence that was in the inner region. After 8 hours, morphological changes in the follicle precluded accurate measurements, so only the first 8 hours after LH were analyzed.

## Results

### Generation of the *Lhr*-COIN mouse line, in which the *Lhr* coding sequence is conditionally replaced by eGFP, and breeding to produce *Lhr*^*INV/+*^ and *Lhr*^*INV/INV*^ mice

The *Lhr* conditional knockout mouse line (*Lhr*-COIN) was generated by inserting a **CO**nditional-by-**IN**version (COIN) cassette—an inverted, promoterless gene-trap cassette^27,28^— into intron 1 of the *Lhr* sequence. The cassette contains a rabbit β-globin splice acceptor (SA) followed by sequences for a P2A self-cleaving peptide, eGFP, and a bovine growth hormone (bGH) polyadenylation signal, in frame with the *Lhr* exon 1 coding sequence. The cassette is flanked by pairs of alternately organized heterotypic lox sites (lox2272 and lox5171) in a head-to-head orientation and is placed in the reverse transcriptional orientation relative to *Lhr* (**Figure 1**). Additional information about the generation of this mouse line is provided in the Supplementary Methods^20^. We use the term “*Lhr*” to refer to the gene formally named *Lhcgr* (luteinizing hormone/choriogonadotropin receptor), because mice lack chorionic gonadotropin, and luteinizing hormone is the sole physiological agonist for this receptor.

**Figure 1.**
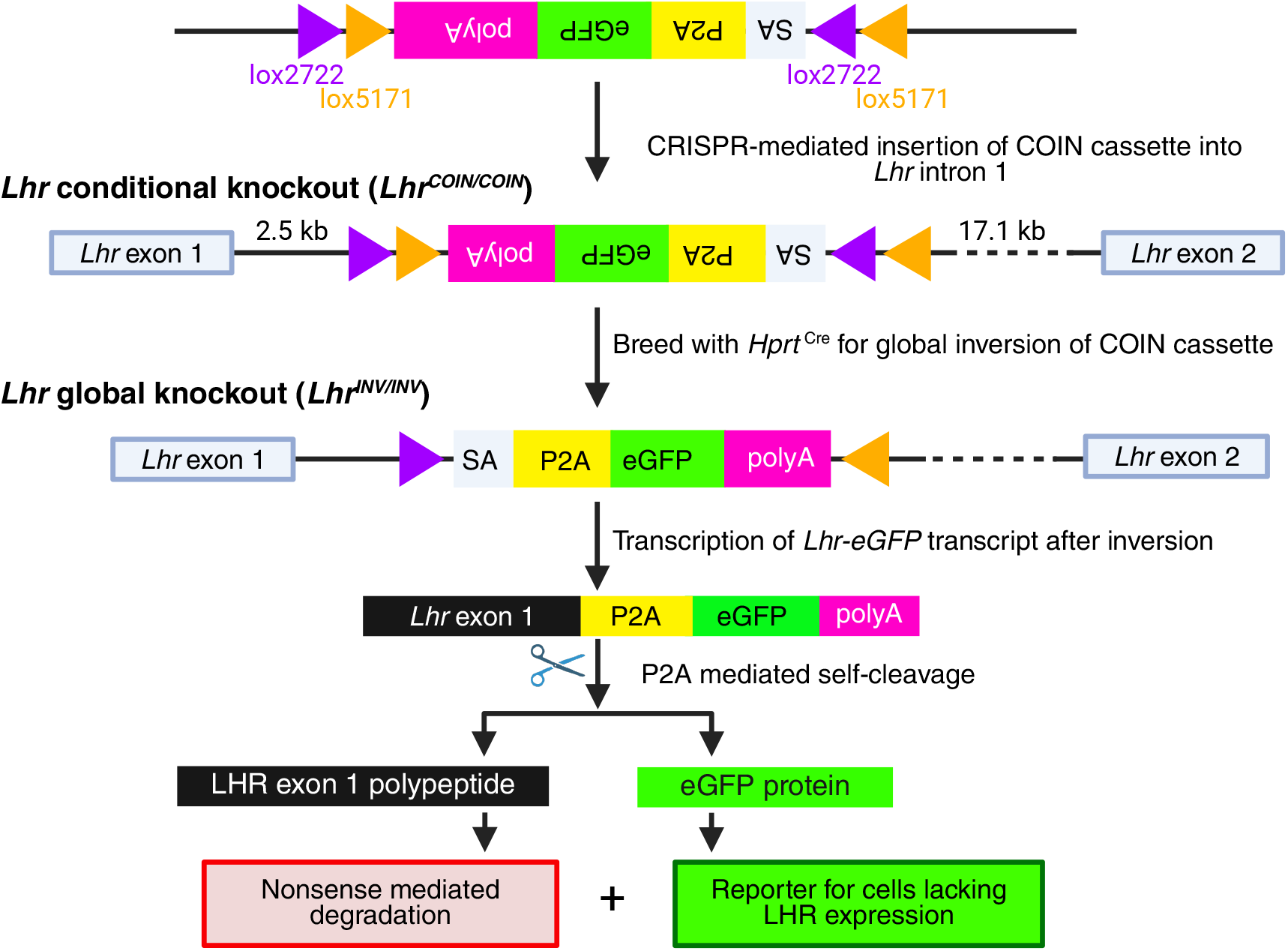
Design of the *Lhr*-COIN conditional knockout mouse line. The COIN cassette was inserted into *Lhr* intron 1. *Hprt*-*Cre*-mediated inversion of the cassette generates a global knockout mouse in which the *Lhr* coding sequence is replaced by eGFP. Figure generated in Biorender.com. See **Figure S2A**^20^ for additional information. Figure made in Biorender.com.

In the absence of Cre recombinase, the COIN cassette is spliced out during transcription and does not interfere with normal *Lhr* expression. In the presence of Cre, recombination occurs between one of two pairs of oppositely oriented homotypic lox sites (either lox2272 or lox5171) resulting in inversion of the cassette (**Figure 1**). This inversion also positions a single heterotypic lox site on one end of the cassette and another heterotypic lox sequence in between a homotypic lox pair oriented in the same direction. A second Cre-mediated recombination event then excises the heterotypic lox site, leaving a single lox site behind and placing the cassette in the inverted orientation. Consequently, the eGFP coding sequence is placed in the same transcriptional orientation as *Lhr*. Since only one heterotypic lox sequence is left on both ends of the cassette, this inversion can only occur once.

The *Lhr* gene has 11 exons and encodes 700 amino acids with a signal peptide of 26 amino acids at the N-terminus. After the inversion, a fusion transcript encoding *Lhr* exon 1, followed by an in-frame P2A self-cleaving peptide and eGFP, is produced (**Figure 1**). P2A-mediated self-cleavage separates the truncated LHR polypeptide from eGFP. Because *Lhr* exon 1 encodes only 58 amino acids, the truncated LHR product is expected to undergo nonsense-mediated degradation. Expression of eGFP therefore serves as a fluorescent marker to monitor cells lacking functional LHR.

eGFP is an appropriate reporter for LHR expression in the ovary during the preovulatory period, because ovarian levels of both the eGFP protein (**Figure S1**^20^ and **Figure 4A** below) and the LHR protein^13^ are approximately constant over the 12-hour period after LH receptor stimulation. For applications using other tissues, similar controls could be performed for the particular biological conditions.

To obtain mice with global inversion of the COIN cassette (*Lhr*^*INV/+*^), we bred *Lhr*^*COIN/+*^ mice with mice expressing Cre recombinase in the egg (*Hprt*^*Cre/+*^) (**Figure 1**). The genotyping strategy and primers to distinguish the COIN and INV alleles are described in **Figure S2** ^20^ and **Table S1**^20^. The *Lhr*^*INV/+*^ mice showed normal fertility (**Figure 2A**), and isolated follicles from these mice resumed meiosis and ovulated normally in response to LH (**Figure 2B,C**). *Lhr*^*INV/+*^ x *Lhr*^*INV/+*^ breeding pairs were used to generate *Lhr*^*INV/INV*^ mice.

**Figure 2.**
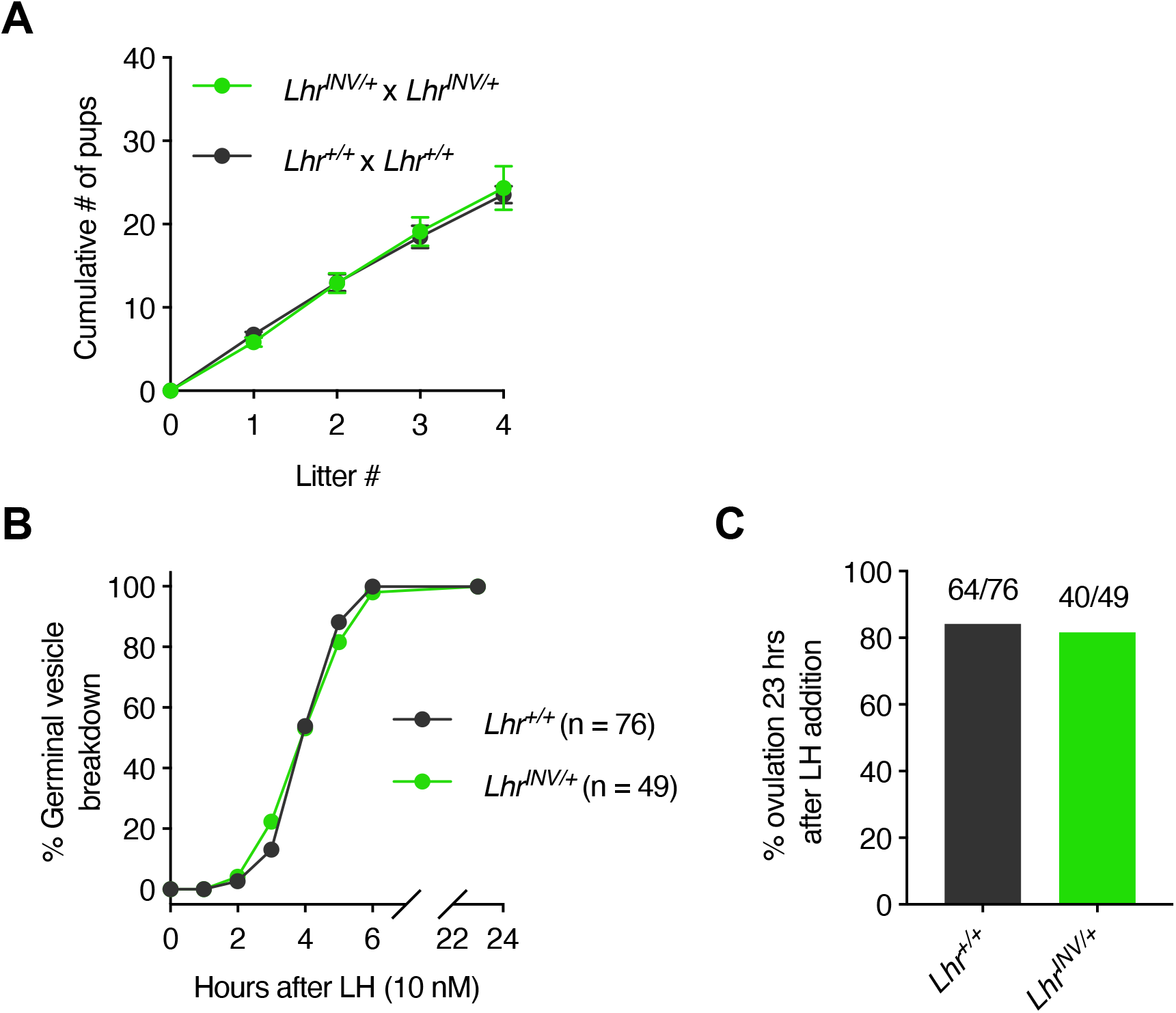
*Lhr*^INV/+^ mice have normal fertility and ovarian follicle responses to LH. **(A)** Cumulative number of pups from *Lhr*^*INV/+*^ x *Lhr*^*INV/+*^ breeding pairs compared to *Lhr*^*+/+*^ x *Lhr*^*INV/+*^ breeding pairs (n = 14 and 11 breeding pairs, respectively; data from the first 4 litters of each; mean ± SEM). **(B**,**C)** Normal LH-induced meiotic resumption and ovulation in *Lhr*^*INV/+*^ mice. B and C show data from 4 experiments with 49 *Lhr*^*INV/+*^ and 76 *Lhr*^*+/+*^ follicles.

The *Lhr* wildtype allele was absent in ovaries from *Lhr*^*INV/INV*^ mice, and no *Lhr* mRNA was detected (**Figure S2C**,**D**^20^). As seen in previous studies of mice lacking the *Lhr*^11,12^, *Lhr*^*INV/INV*^ mice had smaller reproductive tracts (**Figure S3**^20^). In the absence of the *Lhr*, both male and female mice are infertile^11^.

### Ovarian follicles from *Lhr*^*INV/INV*^ can grow to almost full size and respond to FSH but not LH

As previously described for 23-day-old mice with globally deleted *Lhr*^29^, ovaries from 23-26-day old *Lhr*^*INV/INV*^ mice were slightly smaller than those from wildtype mice, with an average weight that was 78% of controls (**Figure S4**^20^). These ovaries contained antral follicles up to ~360 µm in diameter, which is only slightly smaller than those in wildtype ovaries (up to ~400 µm in diameter). The structure of the *Lhr*^*INV/INV*^ follicles was similar to that of wildtype follicles (**Figure 3A**), consistent with a histological section of a *Lhr* global knockout ovary from a previous study^29^.

**Figure 3.**
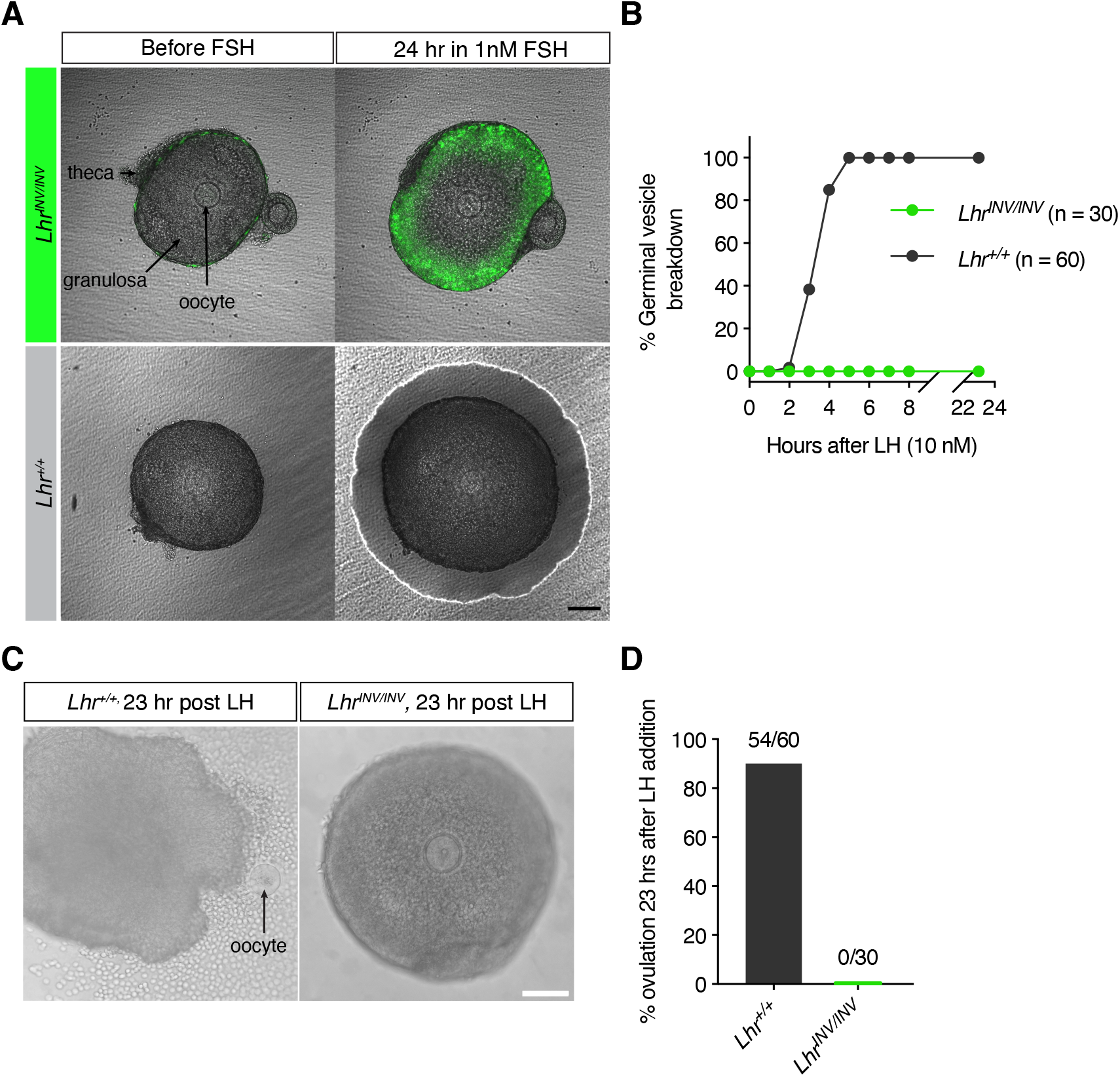
Ovarian follicles from *Lhr*^*INV/INV*^ mice respond to FSH but not LH. **(A)** *Lhr*^*INV/INV*^ and *Lhr*^*+/+*^ follicles, before and after a 24-hour incubation with 1 nM FSH. The scale bar indicates 100 µm and applies to all panels in A. Before applying FSH, eGFP fluorescence was present only in the theca cells (upper left panel). FSH caused expression of eGFP in the outer mural granulosa cells of the *Lhr*^*INV/INV*^ follicles. The white ring around the follicle in the right hand *Lhr*^+/+^ panel is due to the edge of the pool of medium on the top surface of the Millicell. **(B)** Failure of follicles from *Lhr*^*INV/INV*^ mice to resume meiosis in response to LH. Follicles isolated from 23-26-day-old mice were cultured for 24 hours in the presence of 1 nM FSH, then 10 nM LH was added to the dish and follicle-enclosed oocytes were observed hourly to score for GVB. Follicles from *Lhr*^*+/+*^ mice underwent GVB, but those from *Lhr*^*INV/INV*^ mice did not. (**C**) Follicles from *Lhr*^*+/+*^ mice ovulated, releasing the oocyte and some granuluosa cells, but those from *Lhr*^*INV/INV*^ mice did not. The scale bar indicates 100 µm and applies to both panels. (**D**) Failure of follicles from *Lhr*^*INV/INV*^ mice to ovulate in response to LH. Follicles from B were scored for ovulation at 23 hours after LH addition. B and D show data from 4 experiments with 60 *Lhr*^*+/+*^ follicles and 30 *Lhr*^*INV/INV*^ follicles.

Isolated *Lhr*^*INV/INV*^ follicles of 290-360 µm diameter were responsive to FSH (**Figure 3A**) but not to LH (**Figure 3B-D**). FSH stimulates LHR expression in the outer mural granulosa cells of wildtype follicles (see^10^), and in *Lhr*^*INV/INV*^ follicles, it caused eGFP to be expressed in these same cells (**Figure 3A**). The increase in follicular diameter at 24 hours after FSH addition (**Figure 3A**) could be due to stimulation of follicular growth, but could also be affected by time-dependent flattening of the follicle. LH caused oocytes within wildtype, but not *Lhr*^*INV/INV*^ follicles, to resume meiosis, as indicated by breakdown of the large prophase-arrested nucleus known as the germinal vesicle (GVB) (**Figure 3B**). Likewise, LH caused wildtype but not *Lhr*^*INV/INV*^ follicles to ovulate (**Figure 3C,D**).

### LH-induced inward migration of granulosa cells co-expressing the LHR and eGFP, visualized by live imaging

In response to LH, some of the LHR-expressing outer mural granulosa cells in preovulatory follicles migrate inwards towards the cumulus-oocyte complex^13^. This cellular migration has been detected in ovaries fixed at time points between 2 and 12 hours after injecting mice with kisspeptin to induce release of endogenous LH, with amplitude and kinetics similar to a natural LH surge^30^. However, the migration has not been visualized by live imaging. Here we imaged this migratory process in live follicles from mice in which one allele of *Lhr* was replaced by eGFP. The one remaining copy of *Lhr* was sufficient to allow meiotic resumption and ovulation in response to LH (**Figures 2 and 4A**). For imaging of granulosa cell migration, similar results were obtained with *Lhr*^*INV/+*^ mice as described above, or with mice in which the replacement of the *Lhr* with eGFP was made only in the granulosa cells (*Lhr*^*COIN/+*^;*Aro*^*Cre/+*^; see Supplementary Methods^20^)

2-4 follicles (290-360 µm diameter) were dissected from the ovary as a cluster and placed on a Millicell membrane, in the presence of FSH to induce LHR expression (**Figure 4A**). Clusters were used, rather than single follicles, because this minimized flattening of the follicles on the Millicell surface, which allowed better visualization of the LH-induced fluorescence redistribution. Similar results were obtained with clusters of 2, 3, or 4 follicles. eGFP expression was localized to the outer region of the mural granulosa layer, as seen in immunolabelled cryosections of follicles from mice with an HA-tag on the endogenous LHR^10,13^. In the isolated follicle clusters, green fluorescence was visible only on the surface of the follicles that was exposed to the medium (**Figure 4A**), probably due to limited access of FSH to the interior of the cluster, after FSH application in the medium.

**Figure 4.**
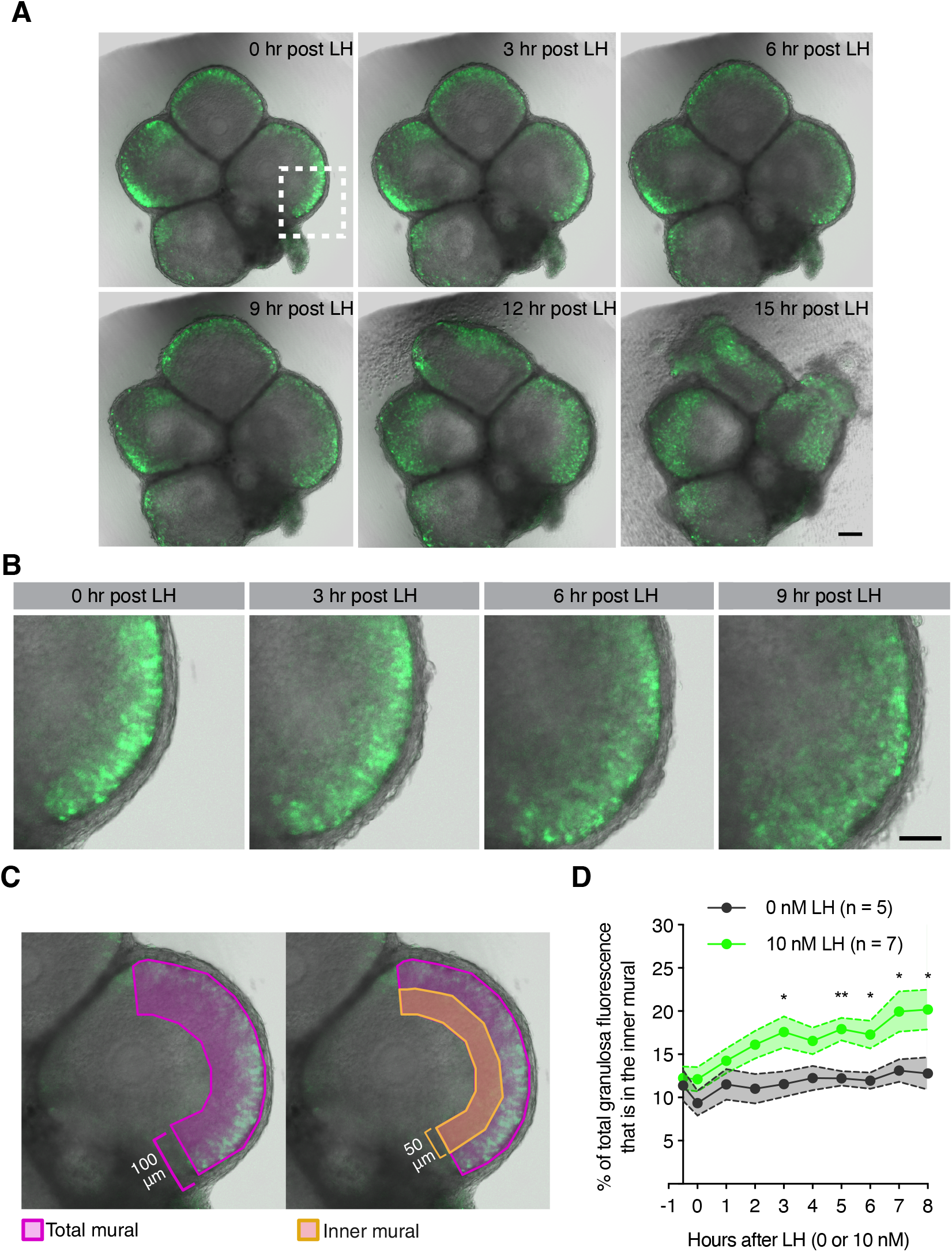
Live imaging of LH-induced ovulation and migration of granulosa cells towards the follicle interior, using follicles expressing one copy of the LH receptor and one copy of eGFP. (**A**) A cluster of preovulatory follicles illustrating LH-induced ovulation. Frames from Supplementary Video 1^20^, at 0-15 hours after applying LH. Scale bar = 100 µm. (**B**) Higher magnification of the region shown by the white dashed box in A, at 0-9 hours after applying LH, illustrating the redistribution of eGFP fluorescence into the inner half of the mural granulosa layer. Scale bar = 50 µm. (**C**) Method for analyzing the redistribution of eGFP fluorescence distribution from the outer to the inner half of the mural granulosa layer, indicative of cell migration. Figure made in Biorender.com. (**D**) Analysis of 7 LH-treated follicles and 5 control follicles, showing the LH-induced relocalization of eGFP. Mean ± SEM; differences between measurements with and without LH were evaluated using a two way ANOVA with the Holm-Sidak correction to compare groups (*p< 0.05; **p< 0.01).

Addition of LH to clusters of preovulatory follicles stimulated ovulation 12-15 hours later (**Figure 4A**, **Supplementary Video S1**^20^). In response to LH, the eGFP fluorescence redistributed towards the follicle interior (**Figure 4A,B**). At 8 hours after applying LH to follicle clusters *ex vivo*, the average percentage of the total mural granulosa layer eGFP fluorescence that was located in the inner half of the mural granulosa layer had increased significantly, from 12% to 20% (**Figure 4C,D**). Controls without LH showed no change in eGFP fluorescence distribution (**Figure 4C,D**, **Supplementary Video S2**^20^). These live imaging results confirm our *in vivo* studies that indicated that LH signaling causes LHR-expressing cells in the outer region of the mural granulosa to move towards the follicle interior^13^.

*Ex vivo* studies using the *Lhr*-COIN mouse line as a fluorescent reporter will facilitate pharmacological studies of the molecular mechanisms and function of the LH-induced granulosa cell migration. Specific application of chemical inhibitors requires follicle isolation, and a fluorescent reporter avoids the multiple steps of fixing, embedding, sectioning, and immunolabelling of follicles for analysis of migration. Inhibitors of interest include those targeting actin regulatory proteins such as cofilin and ARP2/3 that mediate cell migration in other tissues^31,32^.

## Discussion

While the function of the LHR in ovarian mural granulosa cells is well established^1,5–8^, it is unknown if the LHR in theca and interstitial cells might also mediate responses to the mid-cycle LH surge, and the *Lhr*-COIN mice could aid in investigation of this question. The function of the LHR in the theca and interstitial cells in responding to the smaller pulses of LH that are released from the pituitary throughout the female reproductive cell cycle^33^ is also not understood. These questions could be addressed by breeding *Lhr-*COIN mice with mice expressing Cre recombinase in specific ovarian cell types. Cell type-specific inversion of the *Lhr*-COIN cassette might also provide new insights into the function of the LHR in the testis.

In addition, this new mouse line could provide a tool to investigate possible non-gonadal expression and functions of the LHR. Despite evidence for *Lhr* expression in oviduct, uterus, and other female reproductive tissues^18^, female mice with global *Lhr* deletion, when transplanted with pieces of wildtype ovary, reproduce normally, indicating no required role for the LHR in these non-gonadal tissues^29^. However, further investigations of the localization and function of the LHR in cells of these tissues using the *Lhr*-COIN mice could be informative. *Lhr* is also expressed in hematopoetic stem cells, and hematopoesis is elevated in mice with global deletion of *Lhr*^16^. Studies of LHR function in hematopoetic stem cells could be facilitated by use of the *Lhr*-COIN mice. More generally, the *Lhr*-COIN mice may be useful for investigating the expression of the LHR in other tissues and resolving controversies about their possible function^15^.

## Acknowledgments

We thank Jeremy Egbert, Iris Nakashima, Rachael Norris, and Tracy Uliasz for their help with experiments and for valuable discussions. We thank John Eppig for his advice on the manuscript.

## Data availability

The *Lhr*-COIN mouse line has been deposited at the Mutant Mouse Resource and Research Center at The Jackson Laboratory (Bar Harbor, ME) (MMRRC #076250, C57BL/6J-*Lhcgr*^*em2Laj*^/Mmjax, JAX stock #041561). Datasets generated by this study are available by request to the corresponding authors. Supplementary data can be found in the FigShare Repository: https://figshare.com/articles/figure/Supplemental_Data_for_Conditional_replacement_of_the_mouse_LH_receptor_with_GFP_enabling_imaging_of_cell_migration_during_ovulation_Owen_C_M_Lowther_K_M_Kaback_D_Jaffe_L_A_Yee_S_P_/32030100

