## Supplemental Data for "Conditional replacement of the mouse LH receptor with GFP, enabling imaging of cell migration during ovulation"

**Supplementary methods**

**Supplementary figures (S1, S2, S3, S4)**

**Supplementary videos (S1, S2)**

**Supplementary table (S1)**

**Supplementary methods**

**Generation of *Lhr*-COIN mice**

*Lhr*-COIN mice were generated by insertion of a **CO**nditional-by-**IN**version (COIN) cassette into intron 1 of *Lhr* (**Figure 1**). The COIN cassette was synthesized by Genewiz (South Plainfield, NJ). The donor vector was prepared using conventional PCR and molecular cloning to place the 1 kb and 1.1 kb 5’ and 3’ homology arms, respectively.

Two CRISPR sites with high score and low off-targets were identified within intron 1 of *Lhr*, using the CRISPOR database^1^ (https://crispor.gi.ucsc.edu/): LHR CRT, 5’- GCA AAG GAC GGG CAA GCG GA, and LHR CRB, 5’- ATG ATA GGG ATG TCA GGC TG. We tested the efficiency of each sgRNA by electroporation of CRISPR/Cas9 sgRNA ribonucleoprotein (IDT USA) into 20-40 one-cell embryos, which were then cultured to blastocysts. We pooled 2 blastocysts together for DNA extraction, followed by PCR/sequencing to identify the presence of a small insert or deletion (INDEL). The guide test was considered successful when ≥50% of the pooled 2-blastocyst samples contained INDEL at the targeted site as revealed by PCR/sequencing.

The donor vector (20 μl/μg) and sgRNA (25 μl/μg) were co-injected into the pronucleus of one-cell C57BL/6J embryos. Surviving embryos were transferred into pseudopregnant females for subsequent development. Weaned pups were ear notched for PCR genotyping (**Figure S2A,B; Table S1**). Positive founders were further confirmed by PCR/sequencing.

**Generation of mice in which one copy of *Lhr* in the granulosa cells is replaced by eGFP (*Lhr^COIN/+^*;*Aro^Cre/+^* )**

Mice in which one copy of Lhr in the granulosa cells is replaced by eGFP were generated by use of a mouse line expressing Cre recombinase only in the granulosa cells and not elsewhere in the ovary. For this purpose, we bred the *Lhr^COIN/COIN^* mice to mice in which the coding sequence for Cre recombinase was inserted into the 3' UTR of the endogenous aromatase (*Cyp19a*) locus^2^, referred to here as *Aro^Cre^*. Within the ovary, this caused inversion of the COIN cassette specifically in granulosa cells, the only ovarian cell type that expresses aromatase.

1.	Concordet JP, Haeussler M. CRISPOR: intuitive guide selection for CRISPR/Cas9 genome editing experiments and screens. *Nucleic Acids Res*. 2018;46(W1):W242-W245. doi:10.1093/nar/gky354

2.	Unger EK, Burke KJ, Yang CF, Bender KJ, Fuller PM, Shah NM. Medial amygdalar aromatase neurons regulate aggression in both sexes. *Cell Rep*. 2015;10(4):453-462. doi:10.1016/j.celrep.2014.12.040

**Figure S1.** Fluorescence intensity of eGFP in preovulatory follicles expressing one copy of LHR and one copy of eGFP is approximately constant over the 12-hour period following LH addition. Total eGFP fluorescence intensity was measured from time lapse recordings from each of 7 follicles, each similar to that shown in Figure 4A. Fluorescence intensity was determined by tracing the basal lamina and measuring the mean fluorescence within using FIJI. Background fluorescence was measured using wildtype follicles and subtracted. The percent change in fluorescence intensity at each time point was calculated relative to the intensity from an image taken 3-5 minutes after LH addition ("0 hours after LH"). Values indicate mean ± SEM for the 7 follicles.

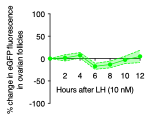

**Figure S2.** Genotyping strategy, and confirmation that *Lhr* DNA and mRNA are absent in ovaries from *Lhr^INV/INV^* mice. (**A**) Genotyping strategy to distinguish *Lhr^COIN^* and *Lhr^INV^* alleles. The COIN and INV alleles were genotyped using two primer pairs, each specific to the 5’ and 3’ regions of the cassette. For the COIN allele, forward primer LHR I1F (intron 1) is paired with reverse primer BGHpAr, specific to the bovine growth hormone (bGH) polyadenylation signal for the 5’ reaction and forward primer eGFPrf is paired with reverse primer LHR I1R (intron 1) for the 3’ reaction. After Cre-mediated recombination, the sequence between the loxP sites inverts to produce the INV allele and the BGHpAr and eGFPrf primers change directionality. To genotype the INV allele, forward primer LHR I1F is paired with eGFPrf and forward primer BGHpAr is paired with LHR I1R to amplify the 5’ and 3’ regions of the cassette, respectively. (**B-D**) PCR analysis of *Lhr* genomic DNA and mRNA in ovaries from *Lhr^INV/INV^*, *Lhr^INV/+^*, and *Lhr*^+/+^ mice. DNA and RNA were isolated using an AllPrep DNA/RNA Mini kit (Qiagen #80204). (**B**) Analysis of *Lhr* genomic DNA in ovaries. Primers LHR I1F and LHR I1R amplify a 308 bp fragment for the wildtype reaction, and primers LHR I1F and eGFPrf amplify a 427 bp fragment for the INV reaction. +/+ mice are positive for only the wildtype reaction, INV/+ mice are positive for both wildtype and INV reactions, and INV/INV mice are positive for only the INV reaction. (**C,D**) Analysis of *Lhr* mRNA in ovaries. The wildtype transcript was detected using forward primer LHR E1-2F (exon1/2 junction) and reverse primer LHR E5R (exon 5), which amplify a 279 bp fragment. The INV allele was detected using forward primer LHR E1F (exon 1) and reverse primer eGFPrf, which amplify a 229 bp fragment. The wildtype transcript was not detected in ovaries of *Lhr^INV/INV^* mice. Figure generated in Biorender.com. Primers are listed in Table S1.

**Figure S2.**

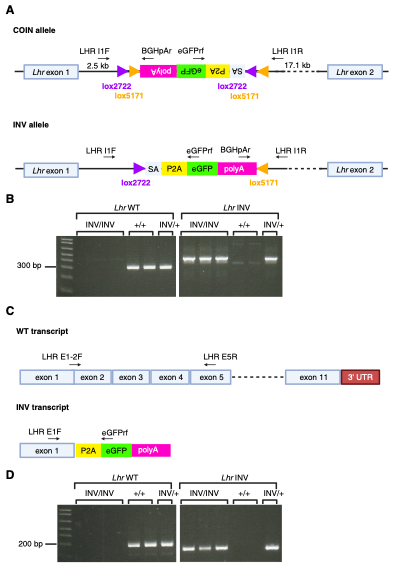

**Figure S3.** *Lhr^INV/INV^* mice have smaller reproductive tracts. Reproductive tracts from 5-month old *Lhr^COIN/COIN^* and *Lhr^INV/INV^* females **(A)** and males **(B)**. Scale bars = 1 cm.

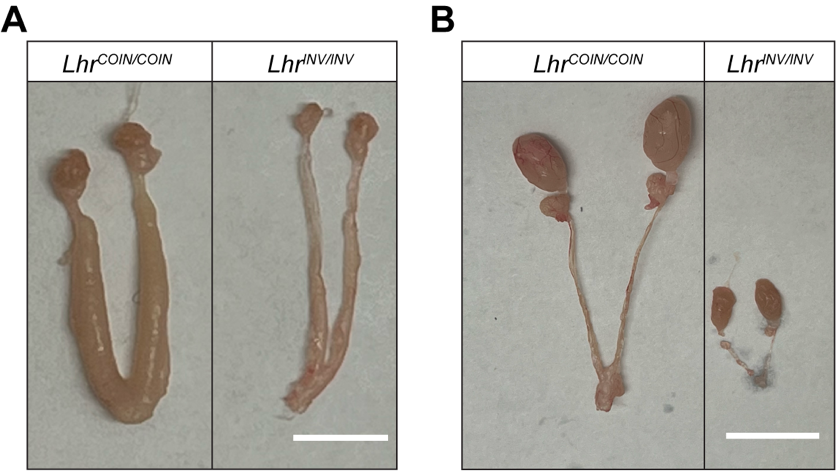

**Figure S4.** Weights of ovaries from 23-26 day old wildtype and *Lhr^INV/INV^* mice. Ovaries from 3 mice of each genotype. Mean ± SEM. Data were analyzed via an unpaired t-test (*p < 0.05).

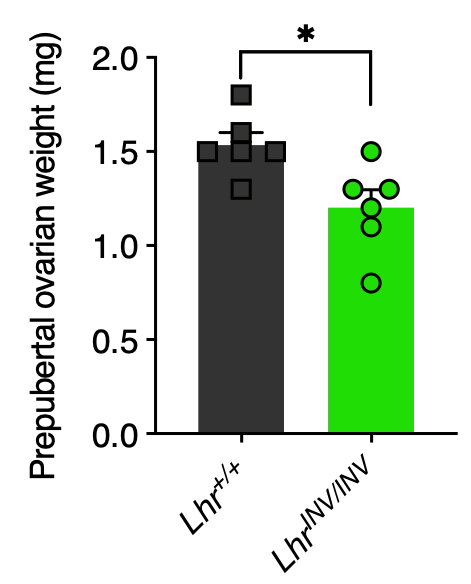

**Supplementary Video S1.** Time lapse imaging of a cluster of preovulatory follicles for a period of 15 hours after adding LH, showing LH-induced inward migration of granulosa cells expressing LH receptors and eGFP, and ovulation.

**Supplementary Video S2.** Time lapse imaging of a control cluster of preovulatory follicles incubated for 15 hours without LH, showing no change in localization of granulosa cells expressing LH receptors and eGFP, and no ovulation.

**Supplementary Table S1.** Primers for distinguishing *Lhr* COIN and INV alleles and transcripts.

| **Target region** | **Primer Name** | **Primer sequence (5’ to 3’)** | **Size of amplicon** |
| --- | --- | --- | --- |
| 5’ COIN allele | LHR I1F | GGAGAATGACAGGCAGAGGGC | 370 bp |
|  | BGHpAr | GCTGGGGATGCGGTGGGCTC |  |
| 3’ COIN allele | eGFPrf | CAGCTCCTCGCCCTTGCTCACC | 413 bp |
|  | LHR I1R | CATGATCCATTTTGCTGCCTGACG |  |
| 5’ INV allele | LHR I1F | GGAGAATGACAGGCAGAGGGC | 427 bp |
|  | eGFPrf | CAGCTCCTCGCCCTTGCTCACC |  |
| 3’ INV allele | BGHpAr | GCTGGGGATGCGGTGGGCTC | 274 bp |
|  | LHR I1R | CATGATCCATTTTGCTGCCTGACG |  |
| WT allele | LHR I1F | GGAGAATGACAGGCAGAGGGC | 308 bp |
|  | LHR I1R | CATGATCCATTTTGCTGCCTGACG |  |
| INV transcript | LHR E1F | GTGCTGGCAATGCTGGTGCT | 229 bp |
|  | eGFPrf | CAGCTCCTCGCCCTTGCTCACC |  |
| WT transcript | LHR E1-2F | GGCCTCGCCCGACTATCTCTC | 279 bp |
|  | LHR E5R | CGAAACATCTGGGAGGGTCCGG |  |
